# Clearing method adapted to FFPE tissues for 3D imaging of nerve fibers, B cells, and tertiary lymphoid structures

**DOI:** 10.1101/2024.05.05.592575

**Authors:** Safa Azar, Laïla Letaïef, Guy Malkinson, Sabrina Martin, Pierre Validire, Jean-Luc Teillaud, Marie-Caroline Dieu-Nosjean, Isabelle Brunet

**Affiliations:** Center for Interdisciplinary Research in Biology (CIRB), Collège de France, CNRS, INSERM, PSL Research University, Paris, France; Université Antonine - Baabda, Lebanon; Sorbonne Université UMRS1135, Faculté de Santé, Paris, France; Inserm U1135, Paris, France; Immune Microenvironment and Immunotherapy Laboratory, Center of Immunology and Infectious Diseases, CIMI-Paris, Paris, France; AMAL Therapeutics, Geneva, Switzerland; Department of Pathology, Institute Mutualiste Montsouris, Paris, France

**Keywords:** 3D imaging, FFPE biopsy, iDISCO, lymphoid structure, nerve fiber

## Abstract

Biomedical samples are commonly used for histological examination after their inclusion in paraffin (Formalin-Fixed Paraffin-Embedded (FFPE) biopsy). However, they provide minimal information about the interactions between different tissue components such as blood vessels, nerves, and cellular aggregates. Here we present a modified iDISCO tissue-clearing method that we term miDISCO^+^. miDISCO^+^ can be used for analyzing FFPE samples, requires the use of only one single FFPE sample, and allows to acquire a detailed image of the 3D tissue/organ architecture and the relevant cellular interactions. The addition of an antigen retrieval step to the protocol enables to use antibodies whose binding is sensitive to formalin fixation due to antigen masking. This method enabled us to detect CD20^+^ B cell follicles and to show that they are in close contact with TH^+^ sympathetic nerve fibers in FFPE biopsies of human palatine tonsils, and to detect CD20^+^ B cells in lung tumors and observe their 3D organization within tertiary lymphoid structures. Thus, miDISCO^+^ could be a potent tool for clinicians to refine diagnosis and to select optimal personalized treatment by giving an integrated 3D view of the tissue structures and cellular interactions in single FFPE samples.

## Introduction

Simultaneous *in situ* analyses of interconnected systems like neurological, vascular, and immune systems are becoming critical for unveiling hidden cross-talks in a given tissue. While formalin fixation followed by paraffin embedding (FFPE) is the standard method used for diagnosis after surgery and biopsy removal (Holdhoff et al., 2019; Banik et al., 2020), it also exhibits significant limitations in investigating these interconnected networks because FFPE samples are routinely processed by cutting blocks. By contrast, three-dimensional (3D) imaging allows a comprehensive analysis performed at high cellular resolution while preserving tissue structures. 3D imaging techniques that rely on clearing techniques (Ertürk et al., 2012; Renier et al., 2014; Susaki et al., 2014) allow the study of large biological samples such as whole murine organs. These techniques allow high-resolution 3D imaging of specimens from various tissues and organs, of perinatal mouse embryos or adult mouse or rat whole body (Pan et al., 2016) at single-cell resolution. An improved clearing method termed iDISCO^+^ (immunolabeling-enabled 3D imaging of solvent-cleared organs) (Renier et al., 2016) has been also used to construct a collection of 3D data sets of immunolabeled and transparent human embryos and fetus images (https://transparent-human-embryo.com/) (Belle et al., 2017). Its use has made it possible, for example, to show netrin-1-dependent sciatic nerve invasion by blood vessels during embryogenesis and its decrease during the early post-natal period (Taïb et al., 2022). A protocol for nanobody (V_H_H)-based whole-body immunolabeling and clearing method (termed vDISCO) that renders whole mice transparent in three weeks, enabling the analysis of an entire body at cellular resolution, has been also recently described (Cai et al., 2023).

3D imaging that exhibits a >10-fold imaging depth as compared to intravital imaging has also the remarkable benefit of disclosing interactions between cancer cells and tumor microenvironment. It allowed, for example, to visualize mouse brain microvasculature and the route of invasion by glioblastoma cells in the tumor core (Lagerweij et al., 2017). 3D light-sheet fluorescence microscopy (LSFM) of cleared solid tumors also identified phenotypic heterogeneity in angiogenesis and in the epithelial-to-mesenchymal transition at single-cell resolution in whole mouse and human FFPE biopsy samples (Tanaka et al., 2017). Thus, 3D reconstruction allows to perform an integrated analysis of cellular networks within tumors and in surrounding tissues, a major advancement as the tumor microenvironment is highly heterogeneous. Identifying where distinctive cell types and structures are located inside/around the tumor and how the different cellular networks are remodeled during tumor evolution is a major challenge for a better understanding of tumor progression and control. The capacity to perform 3D imaging of FFPE tissues obtained after surgery and biopsy removal is of considerable interest for diagnosis and monitoring *in situ* response to treatments. Thus, we developed a protocol based on the iDISCO method that enables testing of FFPE murine and human samples with antibodies including those requiring antigen retrieval due to formalin fixation. The use of FFPE samples for 3D imaging based on clearing methods followed by LSFM has been rarely described (Tanaka et al., 2017), most of the reports being based on the use of PFA-fixed tissues (Pisano et al., 2022; Cai et al., 2023; Frenkel et al., 2023).

In the present report, entire murine organs and lung tumor samples of patients with non-small cell lung cancer (NSCLC) were 3D-imaged by ultramicroscopy following a modified iDISCO^+^ (miDISCO^+^) method adapted to FFPE samples. miDISCO^+^ enabled us to spatially localize, in single samples, the spatial localization of nerve fibers and immune cells, in addition to the vasculature, in normal tissues (brain, spleen, and kidney) and NSCLC tumor samples. Tertiary Lymphoid Structures (TLS) that develop in solid tumors (Dieu-Nosjean et al., 2008; Germain et al., 2014) and have been associated with a better response to anti-immune checkpoint (ICP) therapies (Helmink et al., 2020), were visualized using miDISCO^+^. The miDISCO^+^ method also allowed us to detect CD20^+^ TLS-B cells in close contact with TH^+^ sympathetic nerve fibers in FFPE human palatine tonsil samples, offering novel practical solutions for a deeper understanding of FFPE pathological-derived samples.

## Results

### Formalin-Fixed Paraffin-Embedded tissues can be 3D-imaged using a modified iDISCO method

First, we examined whether the tissue clearing of FFPE samples can be used for 3D imaging like PFA-fixed samples. The clearing step was based on the iDISCO^+^ protocol (Renier et al., 2016), but optimized to perform 3D immunostaining of FFPE samples (hereafter termed “modified iDISCO^+^, miDISCO^+^). In this miDISCO^+^ protocol, an antigen retrieval step was included. Before applying the optical clearing technique, overnight deparaffinization at 70°C followed by overnight incubation with Clearene solution at room temperature (RT) and ethanol baths were used to guarantee the complete removal of paraffin (**Figure 1A**). At the gross anatomical level, no differences between PFA-fixed and FFPE organs (kidney and lungs) were observed after optical clearing using the miDISCO^+^ protocol (**Figure 1B**). Figure 1B shows also that the FFPE brain after clearing is slightly bigger than the PFA-fixed brain. Globally, no evidence of tissue deterioration, shrinkage, or swelling was detected in fixed tissues in both conditions. Since formalin fixation and subsequent paraffinization procedure compress the texture of the biopsy, incubation times for both primary and secondary antibody labeling were extended by one additional day per antibody, as compared to the iDISCO^+^ protocol, to ensure a better antibody penetration into tissue (**Figure 1A** and see Methods for details).

**Figure 1.**
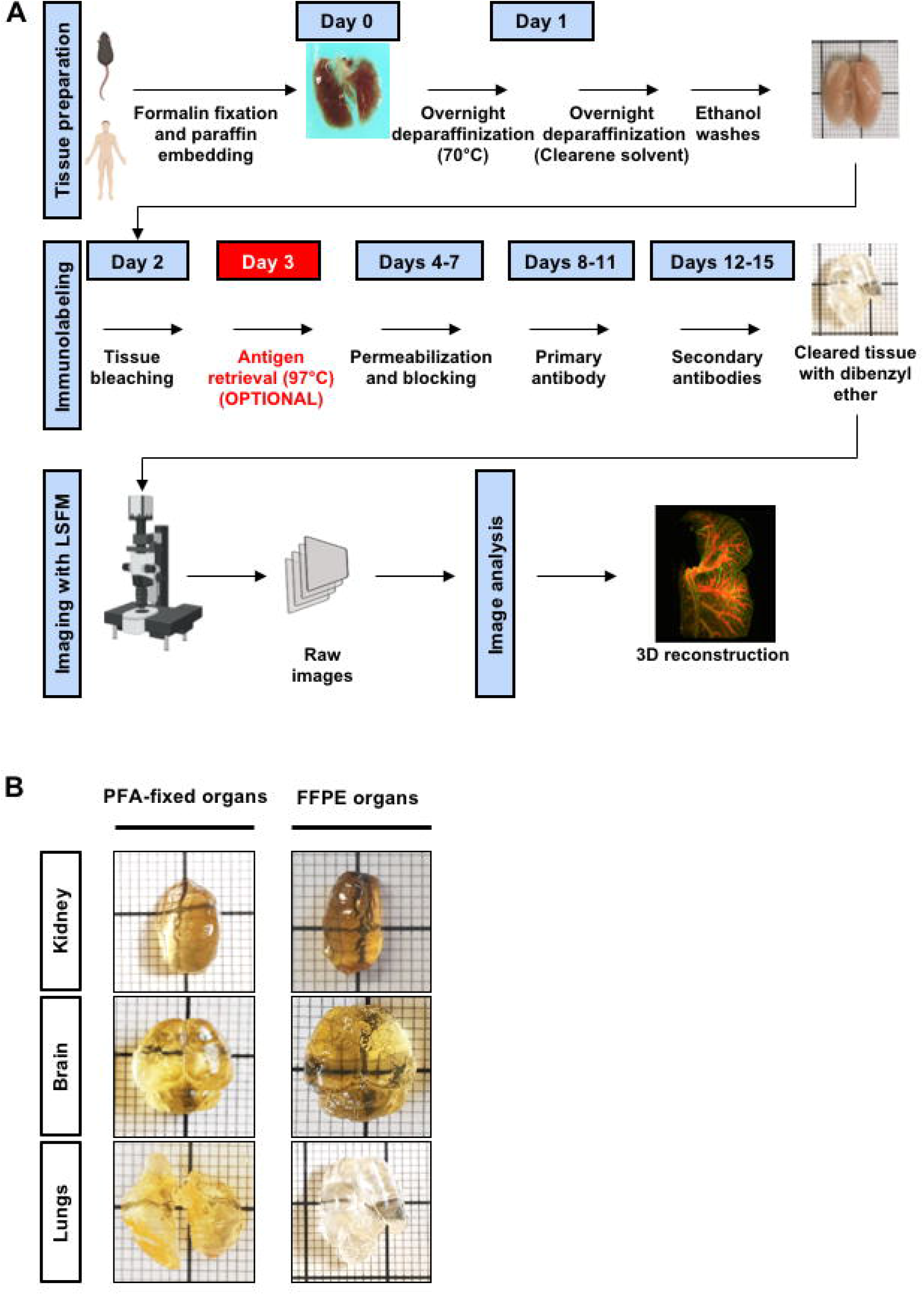
Overview of the clearing and antibody-labeling workflow and comparison of PFA-fixed and FFPE organs after optical clearing with miDISCO^+^. (A) miDISCO^+^ protocol designed for 3D imaging of human and mouse FFPE samples. (B) PFA-fixed and FFPE murine organs (kidney, brain, and lungs) after optical clearing with miDISCO^+^.

We next examined whether clearing paraffin-embedded samples provoked changes in tissue structure and immunostaining characteristics in comparison with miDISCO^+^ cleared PFA-fixed samples. Staining of alpha-smooth muscle actin (α-SMA, a marker of myofibroblasts surrounding blood vessels) was performed in PFA-fixed and FFPE murine kidneys using the miDISCO^+^ protocol, to see if we could detect structural differences in the vascular system between PFA and FFPE. Using LSFM, 3D images were reconstructed. As shown in **Figures 2A** and **2B**, kidney vascularization was intact in both cleared PFA-fixed and FFPE kidney samples (see also **Videos S1** and **S2**). Similar patterns of vascularization were observed (**Figures 2A and 2B**), uncovering the various subdivisions, from renal artery to smaller arterioles, with the same level of detail visible in both conditions. At higher magnification, single smooth muscle cells were visible in FFPE samples (**Figure 2C**, red arrows). The total area and entire volume of the α-SMA immunolabeling were then quantified using Imaris. No significant difference was observed between both types of samples (**Figure 2D**), indicating that both fixation and pre-clearing (removal of paraffin) treatments do not affect the outcome and quality of staining. These results were further confirmed by analyzing another well-described circulatory structure, the circle of Willis, in PFA-fixed and FFPE mouse brains. Arterial anastomosis at the base of the brain was distinguishable regardless of the fixation protocol used (PFA-fixed or FFPE tissue) (**Figure 2E**) (**Videos S3** and **S4**). Qualitatively, the overall structure of the circle of Willis was visible, including posterior communicating arteries and middle cerebral arteries. In addition, antibody penetration depth (at least 400 μm) was similar in both samples (**Figure 2F**) in cerebral cortical tissue.

**Figure 2.**
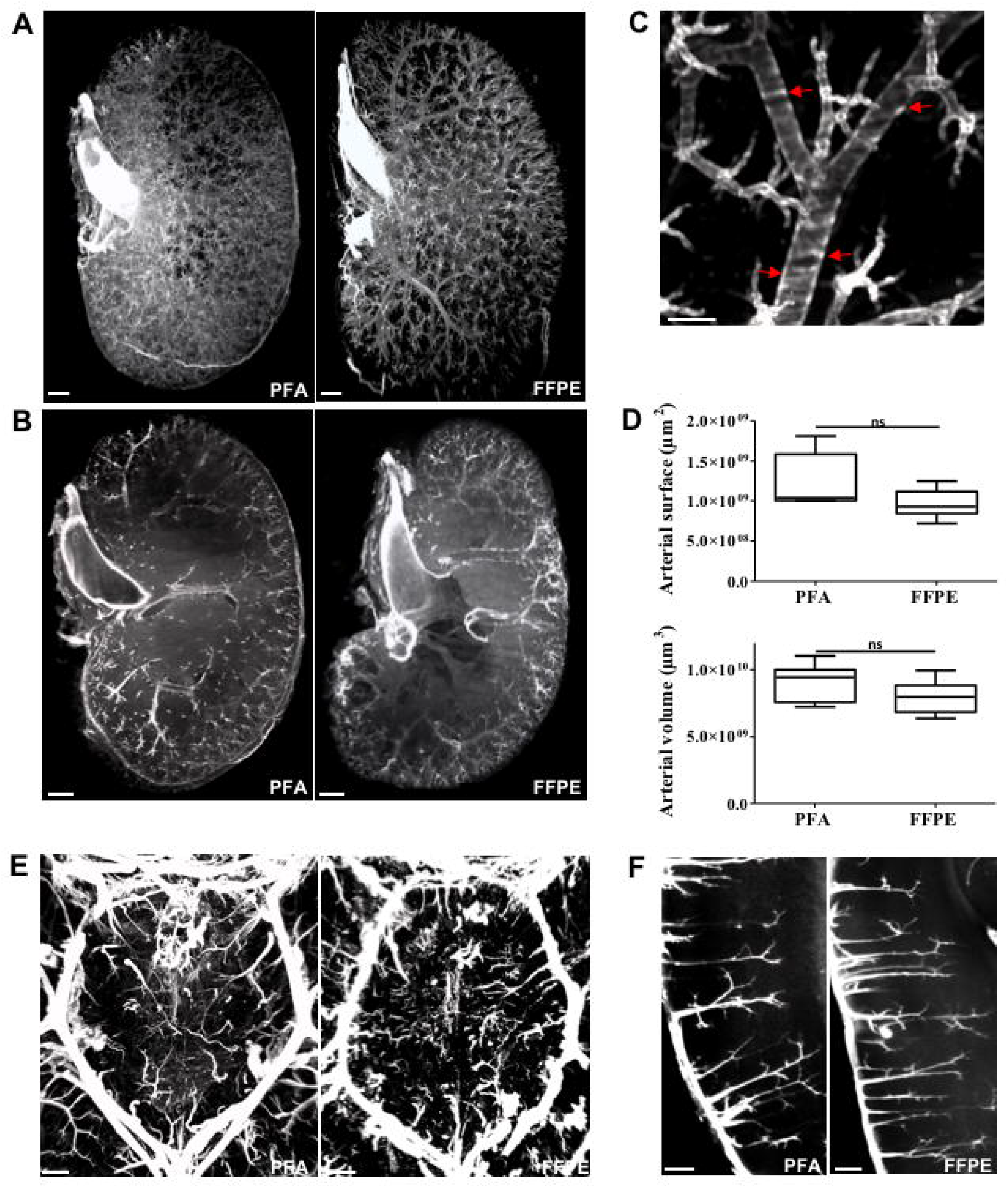
Integrity of vascular structures following tissue clearing of PFA-fixed or FFPE mouse tissues. (A) 3D view of α-SMA staining (shown in white) in PFA-fixed kidney (left panel) and FFPE kidney (right panel). (B) Optical sections at an imaging depth of 600 µm (left panel, PFA-fixed kidney and right panel, FFPE kidney). α-SMA staining is shown in white. (C) Magnification of FFPE kidney showing α-SMA^+^ vessel walls (shown in white). Red arrows point to smooth muscle cells in vessel walls. (D) Comparison plots of α-SMA^+^ areas (upper graph) and volumes (lower graph) between PFA-fixed and FFPE kidneys (PFA-fixed kidneys, n= 8; FFPE kidneys, n= 5). Data are represented as mean ± SEM. ns: not significant. The two-tailed Mann-Whitney test was performed for statistical analysis. (E) 3D imaging of α-SMA staining (white) in the circle of Willis of PFA-fixed (left panel) and FFPE (right panel) mouse brains. (F) Optical sections of the cerebral cortex of PFA-fixed (left panel) and FFPE (right panel) mouse brains. α-SMA staining is shown in white. Scale bars: 500 µm (A, B, E), 100 µm (C), 200 µm (F). Source data are provided as a Source Data file.

Since the technique revealed strong conservation of the general anatomical structures and antigen expression and recognition in paraffin-embedded tissues, FFPE samples were further analyzed in the following 3D imaging experiments.

### miDISCO^+^ method applied to FFPE biopsies allows 3D imaging of subcellular structures and cells labeled with antibodies requiring antigen retrieval

Next, we wanted to evaluate the level of cellular details visible when the miDISCO^+^ method is used on old samples. We thus performed miDISCO^+^ clearing on various human tissue samples that were collected up to 7 years before this study. In this setting, tissue immunolabelling should not exhibit a decreased antigenicity since previous studies have reported stable antigenicity of tissue sections over different periods following embedding (Xie et al., 2011). Thus, the ability to detect subcellular structures was explored using FFPE samples from the human cortex, human tonsils, and from lungs of patients with non-small cell lung cancer (NSCLC). Patient samples had been collected between 4 to 7 years before we tested them using miDISCO^+^. Using our improved protocol, we could detect the Olig2 transcription factor expressed by glial progenitor cells (**Figure 3A**). These cells were distinguishable within the tissue and the method made it possible to discriminate between nuclear and cytoplasmic localizations of the transcription factor (**Figure 3A**, middle and right panels).

**Figure 3.**
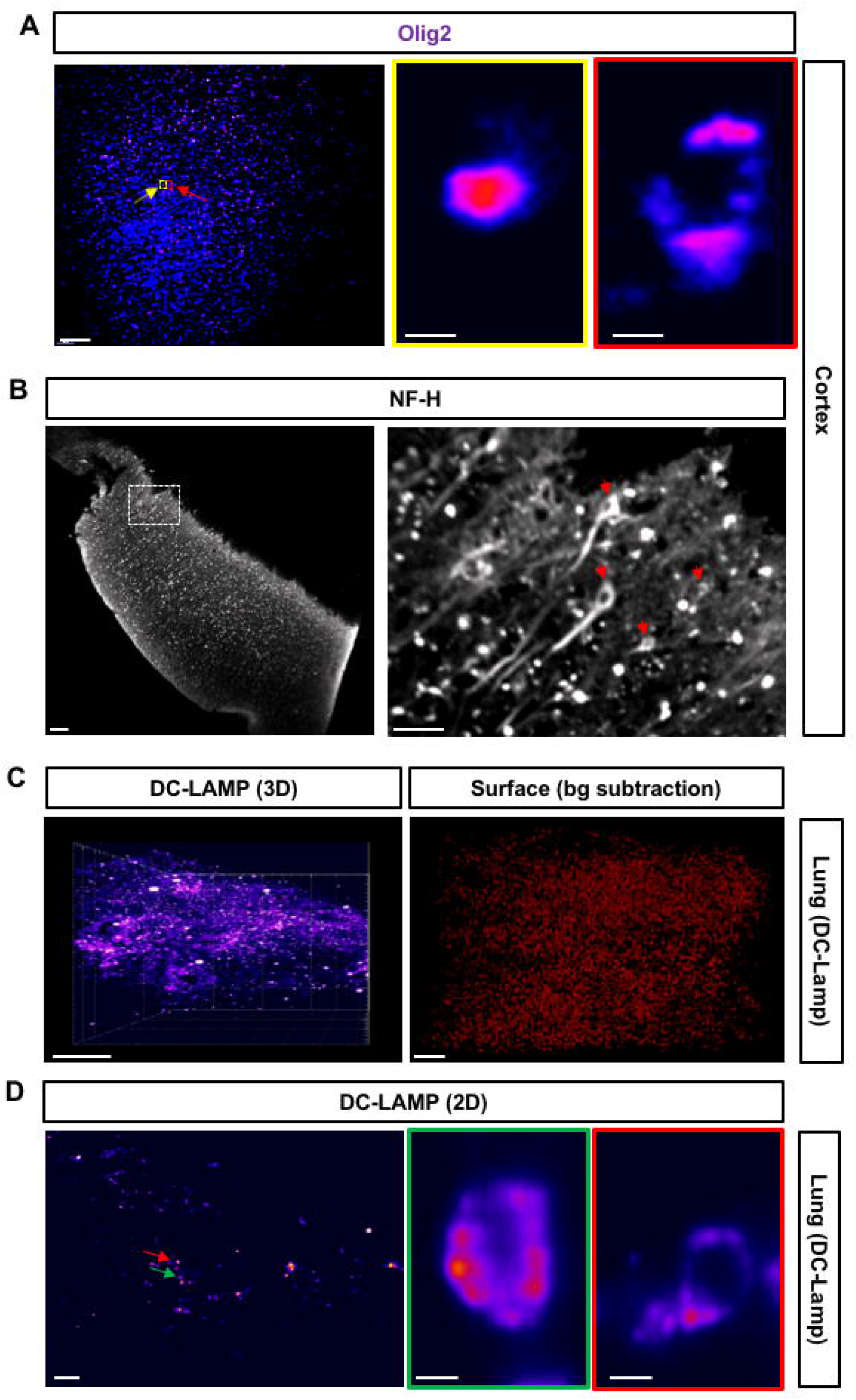
3D visualization of Olig2 transcription factor, NF-H^+^ fibers and DC-LAMP in cleared FFPE human cortex and tonsil. (A) Olig2 immunostaining in the human cortex (displayed as low density in purple to high density in red). The middle and right panels are magnifications of cells observed in the left panel (yellow and red boxed areas indicated by yellow and red arrows). The right panel (framed in red) shows a cytoplasmic localization of Olig2. The middle panel (framed in yellow) shows a nuclear and perinuclear expression of Olig2. (B) 3D image of human cortex NF-H^+^ neurons (shown in white). The right panel is a magnification of the boxed area indicated on the left panel. It shows a distinct profile of NF-H^+^ neuron cell body and neuronal fibers (red arrows). (C) 3D DC-Lamp immunostaining (shown in white) in non-tumoral distant lung. The right panel represents the surface created for background subtraction using a sphericity threshold to filter the noise due to bronchial and vascular empty spaces. (D) 2D DC-Lamp immunostaining obtained after surface reconstruction. Red and green arrows indicate DC-Lamp^+^ cells. The right (framed in red) and middle (framed in green) panels show the cytoplasmic localization of DC-Lamp in the corresponding cells. Scale bars: 50 µm (A, left panel), 5 µm (A, middle and right panels), 150 µm (B, left panel), 50 µm (B, right panel), 100 µm (C), 50 µm (D, left panel), 5 µm (D, middle and right panels).

We were next interested in knowing whether antibodies requiring antigen retrieval for binding to their targets could be used to detect subcellular structures in human tissue samples using miDISCO^+^. We first verified that methanol treatment does not interfere with the antibody labeling of FFPE tissue sections. As shown in **Figures S1A** and **S1B** (left panels), B cells present in human palatine tonsil and lung tumor tissue sections treated with methanol could be efficiently stained by an anti-CD20 antibody, indicating that methanol does not impact anti-CD20 staining efficiency. An antigen retrieval step was then added to the workflow of the miDISCO^+^ method (**Figure 1A**) to test two antibodies whose binding is sensitive to formalin fixation due to antigen masking (**Table S1**). **Figure 3B** (left panel) and **Video S5** show a 3D image of human cortex NF-H^+^ (Neuro-Filament Heavy chain) neurons detected by an anti-NF-H^+^ antibody that required antigen retrieval. Magnification of the boxed area shows the staining of NF-H^+^ neuron cell bodies and fibers (**Figure 3B**, right panel, red arrows). Similarly, **Figure 3C** (left panel) shows that this step makes it possible to detect cells expressing DC-Lamp in non-tumoral distant lungs by 3D imaging using an anti-DC-Lamp antibody that requires antigen retrieval. The surface created for background subtraction using the Imaris sphericity threshold to filter the noise due to bronchial and vascular empty spaces is shown in **Figure 3C** (right panel). DC-Lamp staining was detected at a high resolution within intracellular vesicles of type II pneumocytes in lungs (**Figure 3D**). Remarkably, tissue sections of the human cortex and tonsils that were not subjected to antigen retrieval, treated or untreated with methanol, did not show any positive immunostaining of NF-H or DC-Lamp (**Figures S2A** and **S2B**, right and center-right panels) in contrast to the same tissue sections tested after antigen retrieval (**Figures S2A** and **S2B**, center left and left panels). As already observed with the anti-CD20 staining, immunostaining of NF-H or DC-Lamp treated was not affected by methanol treatment. Of note, the insertion of an antigen retrieval step in the workflow of miDISCO^+^ did not impair the capacity of the selected anti-CD20 antibody that does not require antigen retrieval to stain B cells present in human tonsil and lung tumor tissue sections (**Figure S1A** and **S1B**, upper- and lower-right panels).

Thus, these results demonstrated that the miDISCO^+^ workflow can include an antigen retrieval step if needed and enables a precise detection and localization of subcellular structures and well-defined antigenic targets.

### 3D imaging of B cells, tertiary lymphoid structures and nerve fibers present in human tonsils and lung tumor FFPE samples with the miDISCO^+^ method

These results indicated that miDISCO^+^ can be used to study the localization and spatial relations between cells from various lineages in different tissues using FFPE samples. FFPE lung samples from patients with non-small cell lung cancer (NSCLC) collected 4-7 years before the study were tested. First, immunostaining of α-SMA revealed the vascular patterns of lung tissue of NSCLC patients (**Figure 4A**, left panel, 3D imaging; right panel, 2D optical section). Small arteriole vessels (red arrow) and a large bronchiole (asterisk) could be observed in 2D optical section (**Figure 4A**, right panel). Second, the expression of tyrosine hydroxylase (TH), a marker of sympathetic nerve fibers (Mueller et al., 1969; Lim et al., 2000) that plays a key role in lung function and neuroinflammation responses (Iravani et al., 2002; Straub et al., 2008; Jenei-Lanzl et al., 2015) was then visualized in combination with α-SMA staining. This double labeling showed that thin TH^+^ nerve fibers are close to α-SMA^+^ vasculature, and surrounding arteries (**Figure 4B**, indicated by white arrows in the right panel) as previously shown in studies on the control of sympathetic arterial innervation (Brunet et al., 2014). **Video S6** shows the 3D rendering of anti-SMA and of anti-TH immunolabelling within a FFPE human lung cleared using miDISCO^+^. miDISCO^+^ also allowed to detect CD20^+^ B cells in lung tumors (**Figure 4C**) and to observe their localization and 3D organization in TLS (**Figure 4C**, right panel). FFPE samples from human palatine tonsils that contain large numbers of B cell follicles were also tested to investigate the localization of TH^+^ fibers relative to these lymphoid structures. As shown in **Figures 4D** and **4E** and **Video S7**, TLS-CD20^+^ B cells are in close contact with TH^+^ sympathetic nerve fibers.

**Figure 4.**
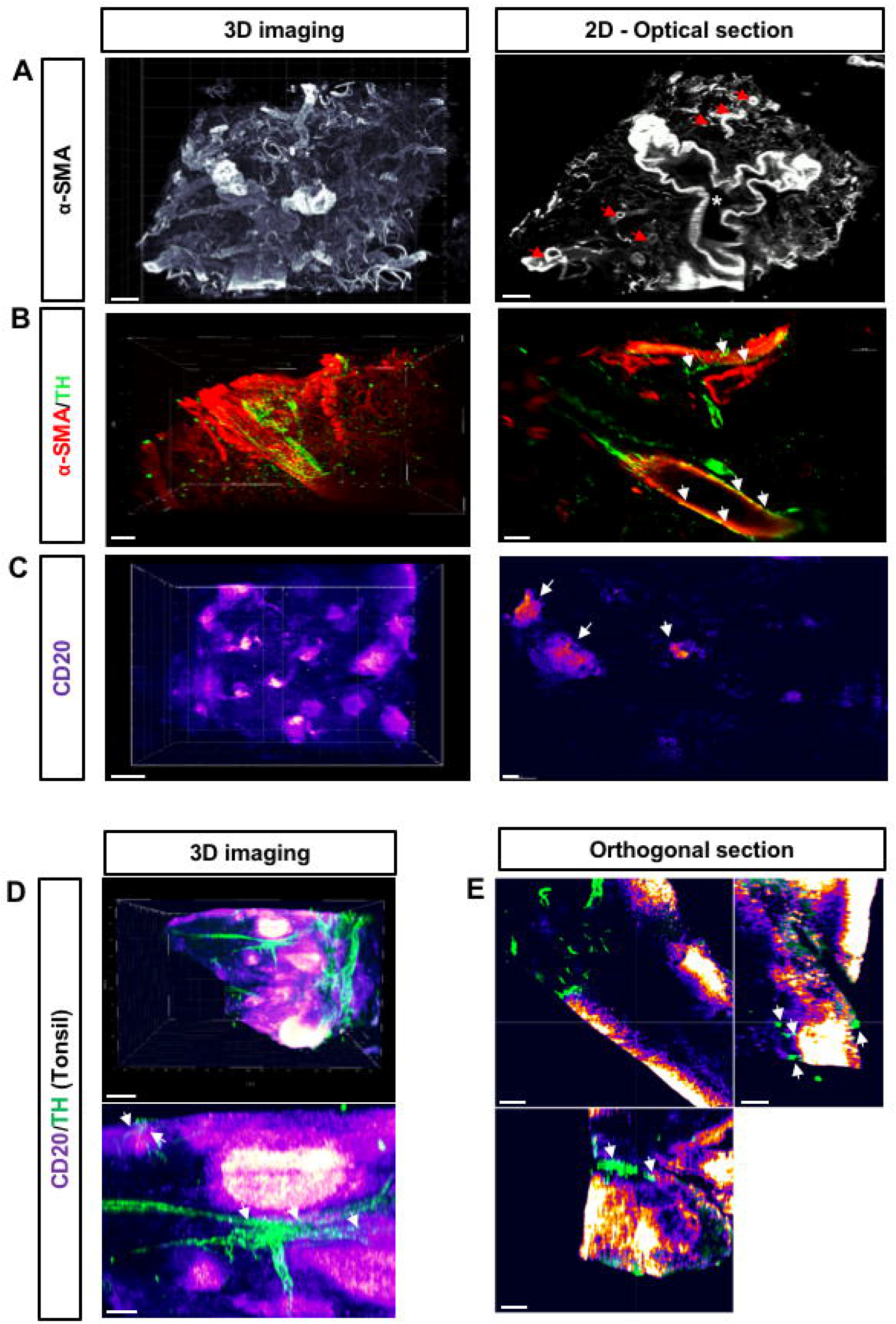
3D imaging of cleared FFPE-fixed human tumor lung specimens from NSCLC patients and 3D view of CD20^+^ B cell lymphoid structures and TH^+^ nerve fibers. 3D and 2D views of lung tumor specimens from NSCLC patients stained with anti-α-SMA-, -TH, and -CD20 antibodies. 2D views are single plans from the 3D imaging. (A) α-SMA staining is represented in white. The asterisk shows a bronchial space delimited with α-SMA staining. Red arrows show vascular staining. (B) α-SMA (red) / TH (green) double staining. White arrows designate TH^+^ fibers in direct contact with α-SMA stained vessels. (C) CD20 staining is represented in pseudo-colors that reflect the intensity of the labeling (from low density in purple to high density in white). TLS-B cells are highlighted by white arrows (right panel). Scale bars: 300 µm (A, B, C, for 3D imaging), 100 µm (A, B, C, 2D for optical sections). (D, E) 3D view of CD20^+^ B cells (from low density in purple to high density in white) and TH^+^ nerve fibers (green) in human tonsils (upper panel). The lower panel (D) and the orthogonal sections (E) show close contacts (white arrows) between CD20^+^ B cell follicles and TH^+^ sympathetic nerve fibers. Scale bars: 300 µm (D, upper panel), 150 µm (D, lower panel, and E).

## Discussion

The present manuscript describes an optimized tissue clearing technique, miDISCO^+^, allowing 3D imaging of formalin-fixed paraffin-embedded tissues. Using this technique, subcellular structures and cells from the nervous and immune systems could be labeled with antibodies requiring antigen retrieval. In addition, CD20^+^ B cells were detected in lung tumors and their 3D organization in TLS could be revealed. Finally, the miDISCO^+^ method allowed us to show that TLS-CD20^+^ B cells are in close contact with TH^+^ sympathetic nerve fibers. 3D imaging of cleared whole tissue has emerged as a potent tool to study the different structures inside organs (Susaki et al., 2015; Vigouroux et al., 2017; Yu et al., 2018) and to unravel 3D cellular networks. It is becoming a necessary method to apply to human tissues, in particular solid tumors, to get an integrated view of their vascular, neuronal, and immune microenvironments. 3D analysis represents a major advantage compared to 2D methods that use tissue sections, notably when cells of interest have a heterogeneous distribution, for example, if they are more abundant in one part of the tissue than in others, and/or when the cutting plane of the tissue makes it difficult to extrapolate to the entire structure of the tissue. The rarity of human tissues for research use and the small sizes of biopsies following surgical tissue resections impose a well-planned use of these samples. The goal of researchers and clinicians is to get as much information as possible from one sample to highlight cell interactions and cross-talk responsible for disease progression. Thus, the access to tissue organization and the ability to spatially relate different structures related to innervation, vascularization, and immune microenvironment at different stages of tumor development represent a formidable tool to understand mechanisms that drive cancer progression and/or therapeutic disease control. However, not all clearing methods can be used successfully for imaging. The nature of organs and tumors being investigated is important. In one study, the CLARITY clearing method resulted in damaged tissue integrity of the murine liver. By contrast, liver samples could be rendered optically transparent using the 3DISCO/iDISCO^+^ methods (Frenkel et al., 2023) and successful immunostaining of intrahepatic microvasculature and tumor cells could be then performed. In the present work, the miDISCO^+^ method permits the detection and localization of CD20^+^ B cells in lung tumors and the observation of their 3D organization in TLS (**Figure 4C**). It also enables to visualize a close contact between TLS-CD20^+^ B cells and TH^+^ sympathetic nerve fibers in FFPE samples from human palatine tonsils (**Figures 4D** and **4E** and **Video S7**). In further experiments, a high-resolution view of nerve fiber/CD20^+^ lymphocytes and their interactions could give a more detailed structural view of these interactions. Our observation is in line with a recent report showing that immune cells present in the direct surroundings of nerve fibers (PGP9.5^+^, a pan-neuronal marker) are mostly CD20^+^ lymphocytes, likely organized in TLS, in pancreatic ductal adenocarcinoma (Heij et al., 2021).

With miDISCO^+^, the possibility of adding an antigen retrieval step further allows us to look for antibodies sensitive to formalin fixation but usable in 3D imaging of tumor microenvironment (TME). Moreover, only one single biopsy is needed when using the method to reveal cellular interactions occurring within TME and possibly to establish a more precise diagnosis, paving the way for optimized clinical treatment in the context of personalized medicine.

Clearing protocols are based on the use of organic solvents that should not affect the general structure of proteins. Protein bonds induced by formalin fixation protect the tissue even after clearing and should ensure an efficient protein extraction, contrary to PFA-fixed tissues. Thus, the miDISCO^+^ method could provide a way to characterize proteins visualized in 3D. Combining 3D imaging methods based on tissue clearing with proteomic and transcriptomic techniques remains a challenge, although important advances have been recently reported (Choi et al., 2021). Using FFPE breast cancer tissue sample slides, a 3D imaging mass cytometry method enabled multiplexed RNA and protein detection, allowing the study of tumor architecture by combining analysis of tissue volumes with single-cell data (Kuett et al., 2022). Moreover, Wang et al. have developed a sequencing-based method (spatially-resolved transcript amplicon readout mapping, or STARmap) for targeted 3D *in situ* transcriptomics in intact tissue allowing to visualize short- and long-range spatial organization of cortical neurons (Wang et al., 2018). The use of multiplexed antibody-based imaging platforms has allowed to obtain high-content imaging data from a wide range of tissues and sample preparations, including FFPE samples, (Hickey et al., 2021), providing direct insights on cell-cell interactions, signaling events, subcellular localizations or translational modifications of proteins, and can complement other assays or single-cell analyses of dissociated tissues.

Our results highlight the proposed workflow (**Figure 1A**) as an essential routine method to unveil information concerning TME, as exemplified herein by the imaging of B cells, lymphoid structures, and nerve fibers in lung tumors. The miDISCO^+^ method should enable the follow-up of interactions between vascular, neurological, and immune networks during tumor progression and cancer therapies. More generally, this method can be used in preclinical studies to assess the impact of drugs targeting one of these networks. We also show that old preserved biopsies could still provide valuable information concerning a given tissue or patient when using miDISCO^+^. This can stand as a powerful tool to extract patient information from previous resections or biopsies (several years ago) and compare them to more recent biopsies from the same patient to study disease evolution, especially at different stages of progression in cancer. The method makes it possible to examine archived human FFPE tissue blocks for immunolabeling and 3D imaging.

As for 2D staining methods, miDISCO^+^ has limitations that include the restricted number of primary antibodies available for FFPE tissue staining compared with the large number of antibodies available for frozen samples. However, more and more antibodies are becoming commercially available for multiplexed imaging, offering the possibility of finding suitable clones usable for miDISCO^+^ 3D imaging. Moreover, common clearing solutions alter emitted light intensity and absorption and emission peaks of Alexa Fluor fluorophores and fluorescent dyes. This increases the risk of unexpected channel crosstalk when multiplexed fluorescence imaging is performed (Eliat et al., 2022). Whether the antigen retrieval step of miDISCO^+^ can be performed only once as opposed to 2D staining techniques remains an open question. This could be the case since it has been recently demonstrated that cleared tissue can be re-embedded into paraffin for further analysis of classical FFPE sections (Tanaka et al., 2017). Thus, all primary antibodies included in multiplex panels should be validated for their ability to still bind relevant epitopes once an antigen retrieval step has been performed. Finally, detailed tissue imaging is currently limited by the low number of fluorescence channels of 3D light-sheet microscopes.

Overall, although miDISCO^+^ exhibits some limitations, it is a very useful method for examining interactions between cellular networks from interconnected systems such as the vascular immune and neurological systems. In particular, the workflow presented in this work represents a useful tool to decipher the 3D architecture of cellular structures within solid FFPE tumors and to follow up its evolution along tumor progression and during cancer therapies. Further improvement will help highlight subtle changes in cellular network cross-talks, particularly in diseases and during disease treatments.

## Acknowledgments

The authors would like to thank Dr. A. Joliot for helpful discussions. We are also grateful to NSCLC patients enrolled in the present study. This study was supported by Institut National de la Santé et de la Recherche Médicale (Inserm) (IB, MCDN, JLT), Sorbonne University, and the Center for Interdisciplinary Research in Biology (CIRB) at Collège de France (IB). The light-sheet microscope used in this work was funded by Leducq Foundation. L.L. was supported first by a fellowship from the French ministère de l’Enseignement supérieur, de la Recherche et de l’Innovation (École doctorale # 394) and then by a fellowship from “La Ligue contre le Cancer”. S.A. was supported by the “Agence Nationale pour la Recherche” #NIRVANA and the Agemed Transversal Program (Inserm).

## Author contributions

S.A, J-L.T., M-C.D., and I.B. conceptualized and designed the study. S.A. and L.L. performed experiments. P.V. provided NSCLC specimens. S.A. and S.M. performed imaging. S.A. and G.M. performed 3D image processing and analysis. S.A., L.L. and I.B., M.-C. D.-N. and J.-L.T. wrote the initial draft of the manuscript. All authors discussed the results and commented on the manuscript text. J.-L.T. edited the final version of the manuscript. I.B. and MCDN provided financial support for the study.

## Methods

### Mouse sample collection

Seven to 14-week-old female C57Bl/6 mice were purchased from Janvier laboratories (Le Genest-Saint-Isle, France) and kept under pathogen-free conditions at the animal facility (UMS 028, School of Medicine, Sorbonne University, Paris, France). All animal studies were performed in compliance with the European guidelines and with the approval of the Charles Darwin Ethics Committee for Animal Experimentation (Paris, France) (agreement N°: A75-13-20). The animals were euthanized and tissues were collected, snap-frozen in liquid nitrogen, and stored at −80°C or fixed in 4 % paraformaldehyde and stored at 4°C. No perfusion-PFA/formalin fixation was performed for mouse tissue in the present work. Other tissues were fixed in formalin and embedded in paraffin according to standard histological procedures (Slaoui and Fiette, 2011). Paraffin blocks were stored at room temperature until analysis.

### Human samples

FFPE non-tumor distant and lung tumor samples were obtained during surgical resection from the Department of Pathology of Institut Mutualiste Montsouris, Paris, France. NSCLC samples were obtained after informed consent. The study was conducted in accordance with the Declaration of Helsinki and its protocol approved by the local ethics and human investigation committee (nos. 2017-A03081-52). Human tonsils and cortex tissues were purchased from Geneticist (Glendale, CA, USA).

### Immunostaining of tissue sections

FFPE human lung, cortex, and tonsil sections of 4 µm-thick were deparaffinized at 70°C in 100% Clearene solvent (Leica Biosystems, Buffalo Grove, IL, USA) solution and washed in ethanol (EtOH) solution followed by methanol (MetOH) dehydration and slide submersion in dH_2_0. Some tissue sections were immersed either in citrate buffer pH 6.0 (Dako, Agilent Technologies, Les Ulis, France; #GV80511-2) [low antigen retrieval (AR) solution] or in EDTA buffer pH 8.0 (Dako; #K800421-2) [high antigen retrieval (AR) solution] for 30 min at 97°C for antigen retrieval. After washing in PBS, tissue sections were then immersed in 100% MetOH solution or PBS for 3 h at room temperature. Slides were then washed in PBS before blocking with Dako protein block (#X090930-2) for 30 min. Primary antibodies (**Table S1**) were added for 1 h at room temperature. The slides were then washed in PBS-0.04% Tween 20 and incubated with revealing antibodies (**Table S2**) for 45 min at room temperature. Nuclei were stained with DAPI (ThermoFisher, Waltham, MA, USA) and scanned with the Axio Scanner (Axio Scan.Z1, Carl Zeiss Microscopy, Jena, Germany).

### Whole-tissue clearing modified iDISCO^+^ method and immunolabeling

For tissue clearing, an iDISCO^+^ protocol (https://idisco.info/idisco-protocol/) was modified and carried out on fresh and FFPE tissue samples. FFPE organs were deparaffinized overnight at 70°C. Then, each tissue was incubated two times in Clearene solvent for 1 h each followed by an additional overnight incubation with Clearene solvent followed by 100% EtOH washes to ensure total removal of paraffin. Freshly recovered organs were fixed in 4% paraformaldehyde (PFA) overnight at 4°C. PFA-fixed and FFPE samples were then dehydrated in serial MetOH solutions (1 h each; 20% MetOH; 40% MetOH; 60% MetOH; 80% MetOH; 100% MetOH), followed by an overnight incubation in 66% dichloromethane (DCM) (Sigma-Aldrich, Saint Louis, MO, USA; #270997) / 33% MetOH solution at room temperature. After 2 washes in 100% MetOH, samples were bleached in chilled fresh 5% hydrogen peroxidase (H_2_O_2_), overnight at 4°C. Samples were then rehydrated in serial MetOH solutions (1 h each; 80% MetOH; 60% MetOH; 40% MetOH 20%) and PBS before immunolabeling. All tissues were incubated with primary antibodies for 4 days, except murine brains that were incubated for 7 days with α-SMA-Cy3 antibody (Sigma-Aldrich; #C6198) (**Table S1**) to ensure a better penetration. For DC-Lamp and NF-H staining, tissues were first incubated in a water bath set at 97°C with citrate buffer pH 6.0 for antigen retrieval (low AR). Samples were then washed 4-5 times for one hour each in sterile PTwH (PBS / 0.2% Tween-20 / 0.01 mg heparin solution). The revealing antibodies (**Table S2**) were then incubated for 4 days. Samples were then transferred to sterile light-protected Eppendorf tubes for washing (10 times) in PTwH (2 mL/sample) at RT and then incubated overnight in PTwH with agitation (Cole-Parmer™ Stuart™ See-Saw Rocker, Thermo Fisher Scientific). To ensure optimum imaging and deposition on the sample holder, tissue samples with small volumes (such as human samples) were then included in 1.5% agarose (Invitrogen, Carlsbad, CA, USA; #16500-500). Tissue clearing was then carried out with MetOH dehydration (as described above) followed by a 3 h incubation with 66% DCM/ 33% MetOH. Next, the samples were washed in 100% DCM (2 x 15 min at RT) and incubated with dibenzyl ether (DBE).

### Tissues imaging

Cleared tissues were then imaged using an LSFM instrument (Ultramicroscope, LaVision BioTec, Bielefeld, Germany) supplied with an optimized macrozoom microscope (MVX-ZB10, Olympus France, Rungis, France). A customized sample holder was generated by LaVision BioTec to allow a better laser sheet exposure of the tissue. For brains, a metal-spiked pocket was used to clench the specimens. The excitation laser lines of 488, 561, and 647 nm were typically used. Z-step size was selected according to the thickness of the laser sheet. 3 µm images were recorded as the z-step size for α-SMA staining of kidneys, tonsils, lungs, and cortex. The z-step size was 4 µm for brain α-SMA staining. Intracellular staining was recorded at 2 µm to ensure better visualization of antibody localization within the cells. Data were collected in a 16-bit TIFF format. Raw data were converted using the Imaris file converter software 9.3.0 (Bitplane, Zurich, Switzerland) into ims. format and images were processed. Videos were created using Imaris 9.3.0 software (Bitplane).

### Blood vessel quantification and Olig2 visualization using Imaris software

Kidney blood vessel areas and volumes were quantified using Imaris software. The surface of α-SMA positive staining was constructed by smoothing the signaling using a median filter. Absolute intensity was chosen for reconstruction excluding all outer signals, thus eliminating all the background. Suitable thresholds were defined to produce the surface rendering. The final surface construction algorithm was saved and applied to the entirety of each data for PFA and FFPE kidneys and statistical values were extracted for both area and surface. The Olig2 surface was created by smoothing the signaling and creating mask. This mask is considered as a valuable function not on the entire image but only in a region of interest (ROI), in this case, Olig2^+^ cells; a filter was then applied to riddle sphericity to subtract any non-cell background caused by the nature of the cortex (unperfused tissue) and the presence of lipofuscin as this cortical tissue was obtained from human [Geneticist].

### Statistical analysis

Values are presented as the mean ± standard error of the mean (SEM). GraphPad 6 (San Diego, CA, USA) was used for statistical analysis. For the comparison of kidney α-SMA areas and volume quantification, the two-tailed Mann–Whitney U-test was used. Kidney samples were encoded for blind imaging and data analysis.

## Resource Availability

### Lead contact

Information and requests for resources and reagents can be directed to and will be fulfilled by the Lead Contact, Isabelle Brunet, PhD.

### Materials availability

This study did not generate new unique reagents.

### Data and code availability

- The data that support the findings of this study will be shared from the lead contact upon reasonable request.
- This manuscript does not report an original code.
- Any additional information required to reanalyze the data reported in this paper is available from the lead contact upon request.

## Supplemental information

**Figure S1. CD20 detection on FFPE sections subjected to methanol baths with an optional antigen retrieval step.** FFPE sections from human palatine tonsils and lung tumor biopsy were deparaffinized and incubated with methanol (MetOH) or not (w/o) allowing to test methanol resistance of the Dako anti-CD20 antibody. An optional antigen retrieval (AR) step was then performed. (A) Human tonsils stained with anti-CD20 antibody (red). (B) Human lung tumor biopsy stained with anti-CD20 antibody (blue). In all panels, the dark blue color in the nucleus shows DAPI staining. Scale bars: 100 µm.

**Figure S2. Comparison of immunostaining of NF-H and DC-Lamp in human cortex and tonsil FFPE tissue sections subjected or not to an antigen retrieval procedure.** FFPE sections from human palatine tonsil and cortex were deparaffinized and incubated with methanol (MetOH) or without (w/o), allowing to test methanol resistance of the Cell Signaling Technology anti-NF-H antibody and the Eurobio Scientific anti-DC-Lamp antibody. Antigen retrieval (AR) was then either performed or skipped to assess the capacity of the antibodies to bind the relevant antigenic targets in miDISCO^+^-cleared tissues. (A) Human cortex stained with anti-NF-H antibody. A strong staining (red) was detectable in tissue sections subjected to antigen retrieval (AR and AR + MetOH). (B) Human palatine tonsil stained with anti-DC-Lamp antibody. Strong staining (white) was detectable in tissue sections subjected to antigen retrieval (AR and AR + MetOH). In all panels, the dark blue color in the nucleus shows DAPI staining. Scale bars represent 20 µm.

**Table S1.** List of primary antibodies used in the study.

**Table S2.** List of secondary antibodies used in the study.

### Electronic supplementary material

**Video S1. Animated 3D rendering of α-SMA labeling in PFA-fixed mouse kidney.** 3D rendering using IMARIS software of α-SMA immunolabelling within a PFA-fixed mouse kidney cleared using the miDISCO^+^ protocol, related to Figures 2A and 2B.

**Video S2. Animated 3D rendering of α-SMA labeling in FFPE mouse kidney.** 3D rendering using IMARIS software of α-SMA immunolabelling within a paraffin-embedded mouse kidney cleared using the miDISCO^+^ protocol, related to Figures 2A and 2B.

**Video S3. Animated 3D rendering of α-SMA labeling in PFA-fixed mouse brain.** 3D rendering using IMARIS software of α-SMA immunolabelling within a PFA-fixed mouse brain cleared using the miDISCO^+^ protocol, related to Figure 2E.

**Video S4. Animated 3D rendering of α-SMA labeling in FFPE mouse brain.** 3D rendering using IMARIS software of α-SMA immunolabelling within a paraffin-embedded mouse brain cleared using miDISCO^+^ protocol, related to Figure 2E.

**Video S5. Animated 3D rendering of α-NF-H labeling in FFPE human cortex.** 3D rendering using IMARIS software of α-NF-H immunolabelling within a paraffin-embedded human cortex cleared using the modified miDISCO^+^ protocol, related to Figure 3B.

**Video S6. Animated 3D rendering of α-SMA and α-TH labeling in FFPE human lung.** 3D rendering using IMARIS software of α-SMA/α-TH immunolabelling within a paraffin-embedded human lung cleared using the miDISCO^+^ protocol, related to Figure 4B.

**Video S7. Animated 3D rendering of α-TH and α-CD20 labeling in FFPE human palatine tonsil.** 3D rendering using Imaris software of α-TH/α-CD20 immunolabelling within a paraffin-embedded human palatine tonsil cleared using the miDISCO^+^ protocol, related to Figure 4D and 4E.

