## Supplementary material for "Clearing method adapted to FFPE tissues for 3D imaging of nerve fibers, B cells, and tertiary lymphoid structures": Table S1

**Table S1.** List of primary antibodies used in the study.

| Primary antibody | Working dilution miDISCO-IF | Antigen retrieval | Species (isotype) | Clone | Manufacturer or Distributor |
|---|---|---|---|---|---|
| α-SMA-Cy3 | 1/200-1/200 | No | Mouse (IgG2a) | 1A4 | Sigma-Aldrich (Saint-Louis, MO, USA) |
| TH | 1/200-1/200 | No | Rabbit | Polyclonal antibodies | Merck-Millipore (Saint Quentin-en-Yvelines, France) |
| CD20 | 1/200-1/70 | No* | Mouse (IgG2a) | L26 | Dako France (Les Ulis, France) |
| NF-H | 1/400-1/400 | Yes | Mouse (IgG1) | RMdO 20 | Cell Signaling Technology (Leiden, The Netherlands) |
| Olig2 | 1/400-NA | No | Rabbit (IgG) | Polyclonal antibodies | IBL-America (Minneapolis, MN, USA) |
| DC-Lamp | 1/100-1:100 | Yes | Rat (IgG2a) | 1010E1.01 | Eurobio Scientific (Les Ulis, France) |

*This antibody can be used for immunostaining of samples previously submitted to an antigen retrieval procedure to test antibodies otherwise unreactive with FFPE tissues.

* This antibody does not require antigen retrieval for binding to $CD20^{+}$ B cells in FFPE tissue sections. It can be also used without any change in immunostaining on samples that have been submitted to antigen retrieval to enable labelling with other antibodies otherwise unreactive with FFPE tissues.
