## Supplementary material for "Clearing method adapted to FFPE tissues for 3D imaging of nerve fibers, B cells, and tertiary lymphoid structures": Table S2

**Table S2.** List of secondary antibodies used in the study.

| miDISCO$^+$ | | | | |
|---|---|---|---|---|
| **Flurochrome** | **Working dilution** | **Species (specificity)** | **Reference** | **Manufacturer or Distributor** |
| Alexa Fluor 647 | 1/400 | Goat (anti-rabbit IgG) | A-21244 | Invitrogen (Carlsbad, CA, USA) |
| Alexa Fluor 555 | 1/400 | Goat (anti-mouse IgG) | A-21422 | Invitrogen (Carlsbad, CA, USA) |
| Alexa Fluor 555 | 1/400 | Donkey (anti-rabbit IgG) | A-31572 | Invitrogen (Carlsbad, CA, USA) |
| Alexa Fluor 555 | 1/400 | Donkey (anti-rat IgG) | Ab15054 | Abcam (Cambridge, UK) |

| Immunofluorescence | | | | |
|---|---|---|---|---|
| **Flurochrome** | **Working dilution** | **Species (specificity)** | **Reference** | **Manufacturer** |
| Alexa Fluor 647 | 1/400 | Chicken (anti-rat IgG) | A-21472 | Invitrogen (Carlsbad, CA, USA) |
| Alexa Fluor 488 | 1/400 | Goat (anti-rat IgG) | A-11006 | ThermoFisher (Courtabœuf, France) |
| Alexa Fluor 647 | 1/400 | Goat (anti-rabbit IgG) | A-27040 | ThermoFisher (Courtabœuf, France) |
| Alexa Fluor 647 | 1/400 | Donkey (anti-mouse IgG) | A-31571 | Invitrogen (Carlsbad, CA, USA) |

* This antibody does not require antigen retrieval for binding to CD20$^+$ B cells in FFPE tissue sections. It can be also used without any change in immunostaining on samples that have been submitted to antigen retrieval to enable labelling with other antibodies otherwise unreactive with FFPE tissues.
