## Supplementary figures and images for "Clearing method adapted to FFPE tissues for 3D imaging of nerve fibers, B cells, and tertiary lymphoid structures"

### Figure S1

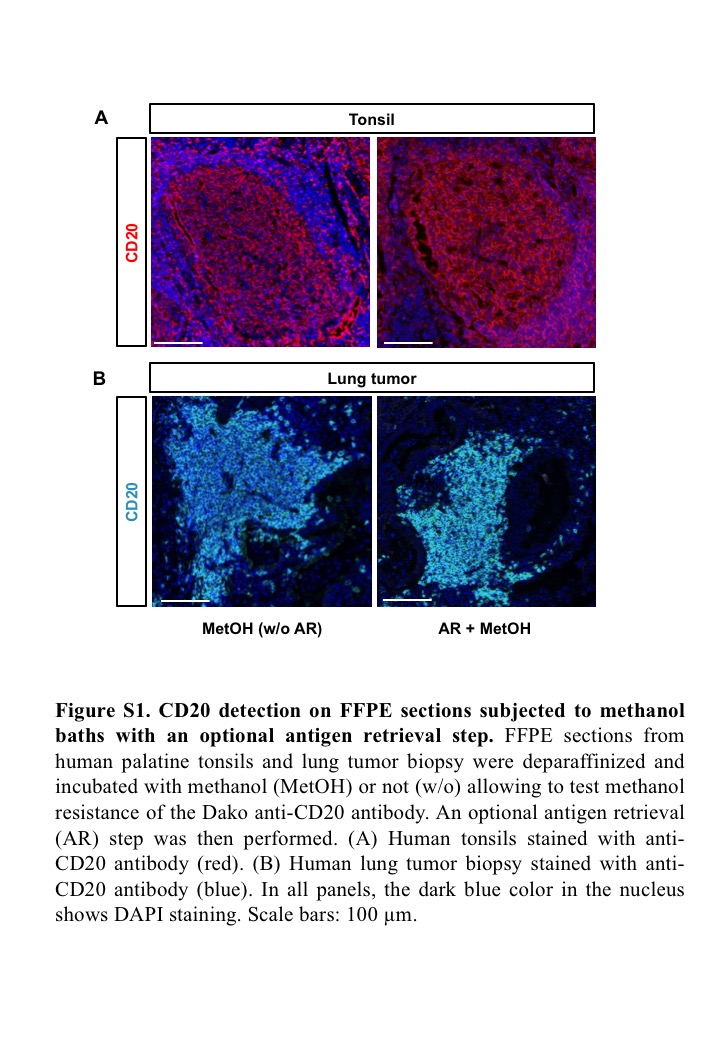

### Figure S2

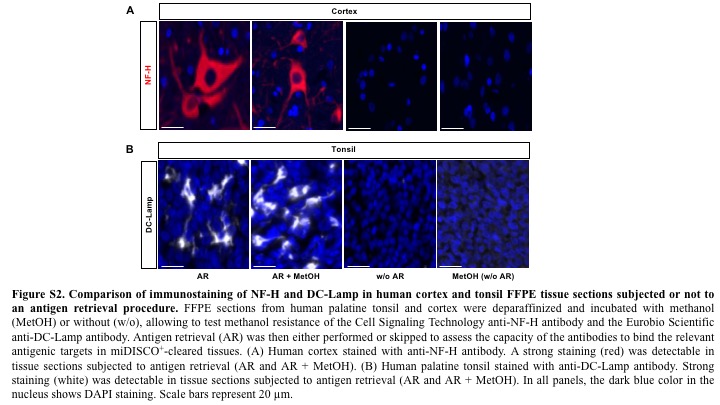
